# RamanBot: Versatile high throughput Raman system

**DOI:** 10.1101/2025.10.03.680377

**Authors:** Khaled Atia, Robert Hunter, Meshach Asare-Werehene, Benjamin K. Tsang, Hanan Anis

**Affiliations:** Department of Electrical and Computer Engineering, University of Ottawa, ON, Canada; Department of Systems Design Engineering, University of Waterloo, ON, Canada; Inflammation and Chronic Disease Program, Ottawa Hospital Research Institute, Ottawa, ON, Canada; Department of Cellular and Molecular Medicine, Faculty of Medicine, University of Ottawa, Ottawa, ON, Canada; Department of Obstetrics & Gynecology, & The Centre for Infection, Immunity and Inflammation (CI3), Faculty of Medicine & Interdisciplinary School of Health Sciences, Faculty of Health Sciences, University of Ottawa, Ottawa, ON, Canada; LifeLabs Medical Laboratory Services, Toronto, ON, Canada

## Abstract

Raman spectroscopy is a powerful tool for qualitative and quantitative analysis in various scientific and industrial fields. However, the development of multisample automated screening remains relatively unexplored. In this paper, we develop RamanBot, a low-cost, easy-to-assemble, and automated Raman spectroscopy system designed for efficient signal collection from samples stored in different types of containers. For the first time, the proposed device introduces the Cartesian motion system, commonly used in 3D printers, to Raman spectroscopy automation. This is achieved by replacing the extrusion head of a commercially available 3D printer with a novel designed “Raman Head”. The Raman head integrates all the necessary optical components required for in-place sample excitation and signal collection. A multimode fiber is used to deliver the excitation laser to the Raman head, whereas the collected Raman signal is delivered to the spectrometer via a fiber bundle. The motion system is programmed to scan predefined sample arrangements using the standard programming language for computer numerical control (G-code). The effect of movement precision on the Raman signal is studied. The introduced device is used in the quantitative analysis of ethanol and methanol. In addition, RamanBot is used to screen six eggs in their commercial packaging with minimal human intervention. The results show that the system is highly stable and capable of delivering reliable Raman measurements, making it a promising solution for high-throughput Raman spectroscopy applications.

## Introduction

Over the years, Raman spectroscopy has emerged as a powerful and versatile analytical technique, finding applications across diverse disciplines such as materials science, life sciences, chemistry, physics, medicine, pharmaceuticals, semiconductor manufacturing, process monitoring, quality control, and forensics [1–5]. The technique operates by measuring the energy shift of incident photons as they interact with molecular vibrations within a sample, enabling both qualitative and quantitative compositional analysis [1, 3]. Despite its broad applicability, Raman spectroscopy has yet to reach its full potential, in part due to the high cost and complexity of developing high-throughput Raman systems [6].

High-throughput Raman screening is particularly important in applications requiring the rapid and reproducible analysis of large sample sets. For example, high-throughput Raman platforms have enabled rapid screening of serum samples for colorectal cancer with approximately 83 % sensitivity and specificity [7], and label-free screening of tens of thousands of eukaryotic cells for biomedical assays [8]. In materials and biomedical contexts, a multifocal Raman spectrophotometer has been used to analyze multiple 3D cell spheroids in parallel—improving throughput by approximately two orders of magnitude [9]. In these scenarios, the ability to collect large datasets with minimal human intervention is critical to accelerate decision-making, reduce operator error, and enable robust statistical analysis.

A significant body of research has focused on integrating Raman spectroscopy with microplate-based platforms, which are widely used in analytical research and medical diagnostics due to their compatibility with multiple detection modalities [7, 10–14]. Standard microplates typically contain 96, 384, or 1536 wells, with capacities ranging from a few nanoliters to several milliliters. However, most Raman systems are designed to accommodate only one or two specific microplate formats, limiting their adaptability. Moreover, many biological and chemical samples are stored in alternative containers, such as capillaries, cuvettes, or centrifuge tubes, which vary in size and configuration [15].

Alternatively, handheld Raman devices [16, 17] offer convenience and portability for point-of-care and on-site measurements. However, despite their widespread adoption, these devices are typically operated manually, which reduces throughput, reproducibility, and measurement consistency, particularly in high-throughput settings or when precise spatial sampling is required [18, 19].

In certain cases, samples must be scanned directly in their original containers to avoid unnecessary handling that could compromise integrity, ensuring more accurate and representative Raman measurements. This diversity of sample formats underscores the need for a flexible, automated Raman system capable of accommodating multiple container types, arbitrary sample arrangements, and direct container-based scanning.

In this work, we leverage recent advancements in motion control systems from fused deposition modeling (FDM) 3D printing to develop a cost-effective, versatile, and high-throughput Raman screening system. Existing automated Raman platforms are often prohibitively expensive, proprietary, or restricted to specific formats, whereas our approach repurposes widely available and inexpensive 3D printer motion systems. This enables a substantial reduction in acquisition time while maintaining high positional precision.

In our system, the extrusion head is replaced with a custom-designed “Raman Head” that houses all optical components necessary for laser excitation and Raman signal collection. Taking advantage of the 3D printer ecosystem, the proposed device is programmable via standard G-code [20, 21]. Pause commands are inserted when the system reaches a sample location to allow the spectrometer to collect a Raman signal. Simultaneously, a synchronized acquisition schedule is passed to the spectrometer to ensure coordination between the motion platform and the spectrometer.

Our approach reduces manual effort, accelerates sample screening, and enables large-scale Raman data collection across a broad range of sample types, bridging the gap between cost-effective instrumentation and high-throughput analytical capabilities. While the current implementation operates with one sample at a time, the idea remains applicable to different excitation configurations, where multiple samples can be excited simultaneously.

The rest of this paper is organized as follows. Section 2 covers the selection of the motion system, the Raman head design, and the overall operation. Section 3 discusses the results of positioning error, and the quantitative analysis of ethanol and methanol in different sample platforms. Section 3 also highlights the system’s adaptability to non-standard sample configurations by scanning commercially purchased eggs in their original carton.

## High-throughput Raman System

### Motion system selection

There are various types of 3D printers available [22]. Among them, FDM printers are the most widely used because of their ease of modification, and the availability of a large support community. The motion system of FDM printers remains an active area of research, with different configurations introduced to optimize speed, precision, and stability [23–26]. The most prevalent configuration is the Cartesian system, valued for its simplicity, reliability, and ease of maintenance. It relies on independent linear movement along the X, Y, and Z axes, offering good stability, resolution, and easy calibration. However, in designs where the Y-axis movement involves shifting the entire print bed, performance is limited by inertia [27]. This limitation is addressed in CoreXY systems [24], where the bed moves only along the Z-axis, while a crossed-belt arrangement drives the print head in the X–Y plane.

Alternatively, Delta printers employ three articulated arms connected to the extrusion head, with the print bed remaining stationary. This kinematic arrangement allows for very high printing speeds and produces a circular build area. However, Delta printers are more complex to calibrate because motion along any axis requires coordinated action of all three motors, and positional accuracy decreases toward the edges of the build plate [28].

Another alternative is the selective compliance assembly Robot Arm (SCARA) systems. It uses a robotic arm that moves in X, Y, and Z, offering a small footprint and mechanical flexibility [29]. However, they are generally slower than Delta designs and harder to calibrate.

The high speed achievable with Delta and SCARA systems is not advantageous in the present work, as it may compromise the stability of the optics. While, the Cartesian configuration provides an optimal balance between accuracy, steadiness, and mechanical simplicity. Most importantly, it sustains cost effectiveness.

For these reasons, we prefer the Cartesian motion system. We chose the open-source Prusa i3 MK3S printer [30], where all the printer pieces are 3D printed, which makes it easy to modify for our purpose. In addition, it offers high mechanical accuracy and strong community support. In our setup, the X–Y motion system of the printer is used to scan samples in different arrangements across the printing bed, while the Z-axis is employed to adjust the focus on each sample. The printer is controlled via standard G-code commands to perform three-dimensional positioning. For example, the command G0 X10 Y10 Z5 F500 moves the head to coordinates (10 mm, 10 mm, 5 mm) at a speed of 500 mm/min.

During acquisition, the motion system is paused at each sampling location, allowing time for the head to come to a complete stop and to mitigate residual vibrations. The spectrometer is programmed to collect a predefined number of spectra per location, with an inter-scan delay accounting for both mechanical travel and stabilization time. This coordinated control ensures high-quality spectral data across all sample positions.

### Raman head

The Raman head is designed after the required optical configuration is selected. In our approach, the excitation light is provided by a 400 mW multimode laser operating at 785 nm, delivered via a 0.22 NA multimode optical fiber entering from the top. The scattered Raman signal is collected from the side through a 0.22 NA seven-core round-to-line fiber bundle (Thorlabs BFL200LS02), as shown in Fig.1:A. This arrangement prevents tangling between the excitation and collection fibers during scanning.

**Fig 1.**
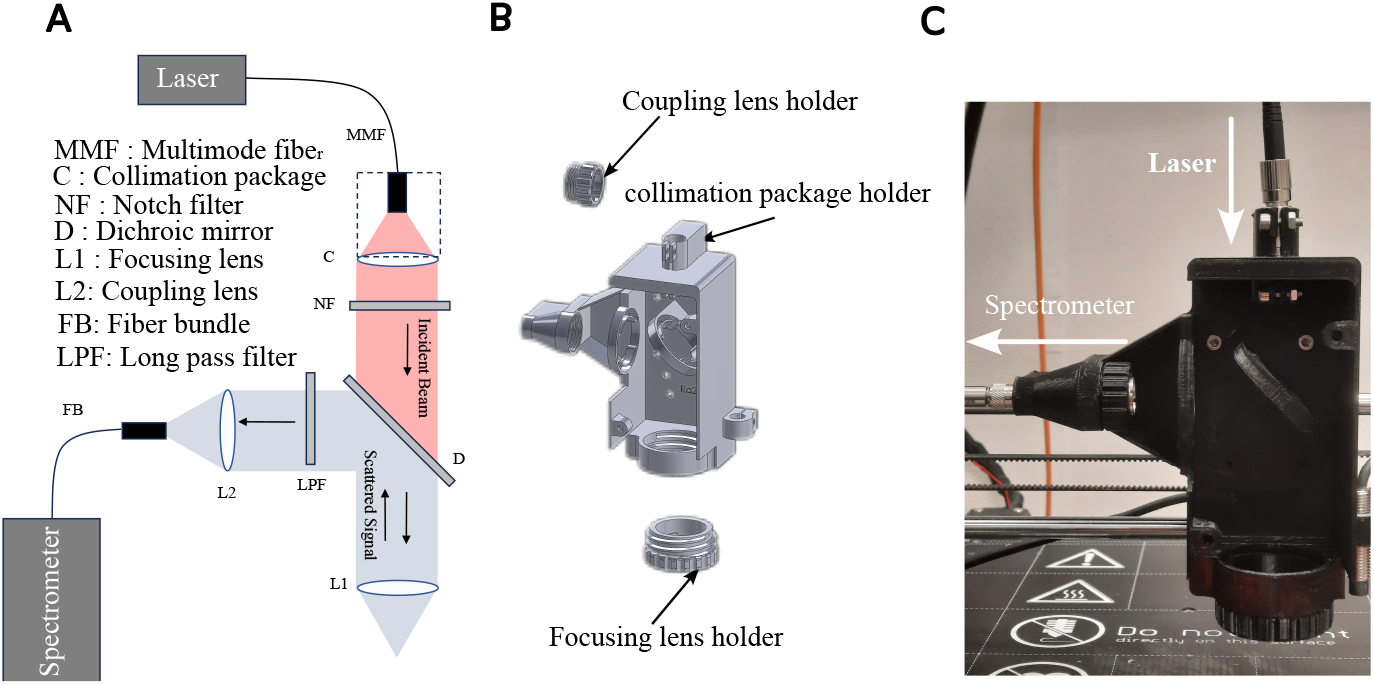
High Throughput Raman. A: Schematic diagram of the optical setup of the Raman head. B: The design of the Raman head that houses all the necessary optics. C: THe 3D printed Raman head after all optics are mounted.

A collimation package (Thorlabs F220APC-780) is used to collimate the output of the multimode fiber that carries the laser light. A 10 nm full-width at half-maximum (FWHM) line filter (Thorlabs FL05780-10) is chosen to suppress broadband background emission of the collimated beam. A short-pass dichroic mirror with a cut-off wavelength of 805 nm (Thorlabs DMSP805) is used to reflect the Raman-shifted light above 805 nm toward the spectrometer side. While, it transmits the 785 nm laser to the objective lens. The laser is then focused onto the sample using a 0.4 NA objective lens.

The reflected Raman signal from the dichroic mirror is passed through a long-pass filter to further attenuate residual excitation light. A 0.22 NA lens is used to couple the Raman signal to the fiber bundle.

Taking into account this optical setup, the original Prusa extrusion head model [30] is redesigned to house these optical components. The new Raman head consists of three main parts as shown in Fig.1:B : (1) the main body, which accepts the multimode fiber input and holds the collimation package together with all optical components except the objective and the coupling lens; (2) a dedicated objective lens holder; and (3) a coupling lens holder for focusing the Raman signal to the fiber bundle. The 3D printed designs can be found in the [31]. The focusing and coupling lens holders were designed as separate components from the main body, allowing lens replacement if needed, without disturbing the integrity of the main structure. They are screwed to the main body for assembly. In the proposed design, we use the flexure of the 3D printed pieces to tightly secure the optical components and maintain precise alignment. The collimation package, however, is additionally fastened with a top screw (Fig. 1:B) to provide enhanced stability, as it is directly coupled to the multimode fiber. Similarly, the fiber bundle is fastened with an SMA adapter with a lock nut (Thorlabs HASMA).

The fully assembled Raman head is shown in Fig.1:C. Although the design is compatible with any spectrometer that accepts a standard fiber input, our experiments use an Andor HoloSpec spectrometer for spectral acquisition [32].

### Overall system operation

Figure 2 shows the overall work flow of the introduced machine. First, samples are placed on the device bed in the desired order. The location of each sample is measured with respect to the Raman head home (0,0,0). The height of each sample is also measured to ensure perfect focus on every sample. The spectrometer acquisition time for each sample is defined with a time margin. The machine pause time at each sample is then defined as the sum of the acquisition time and the time margin. With all these parameters, a sequence of moves separated with pauses at each sample is then converted into a G-code and passed to the machine through pronterface [33], which is open-source software based on Python for controlling 3D printers. It communicates with the printer via USB. Simultaneously, an acquisition schedule is created based on the time taken by the machine to reach a certain sample and the specified acquisition time. The acquisition schedule is then passed to the spectrometer via Solis software [34], which controls our Andor HoloSpec spectrometer [32].

**Fig 2.**
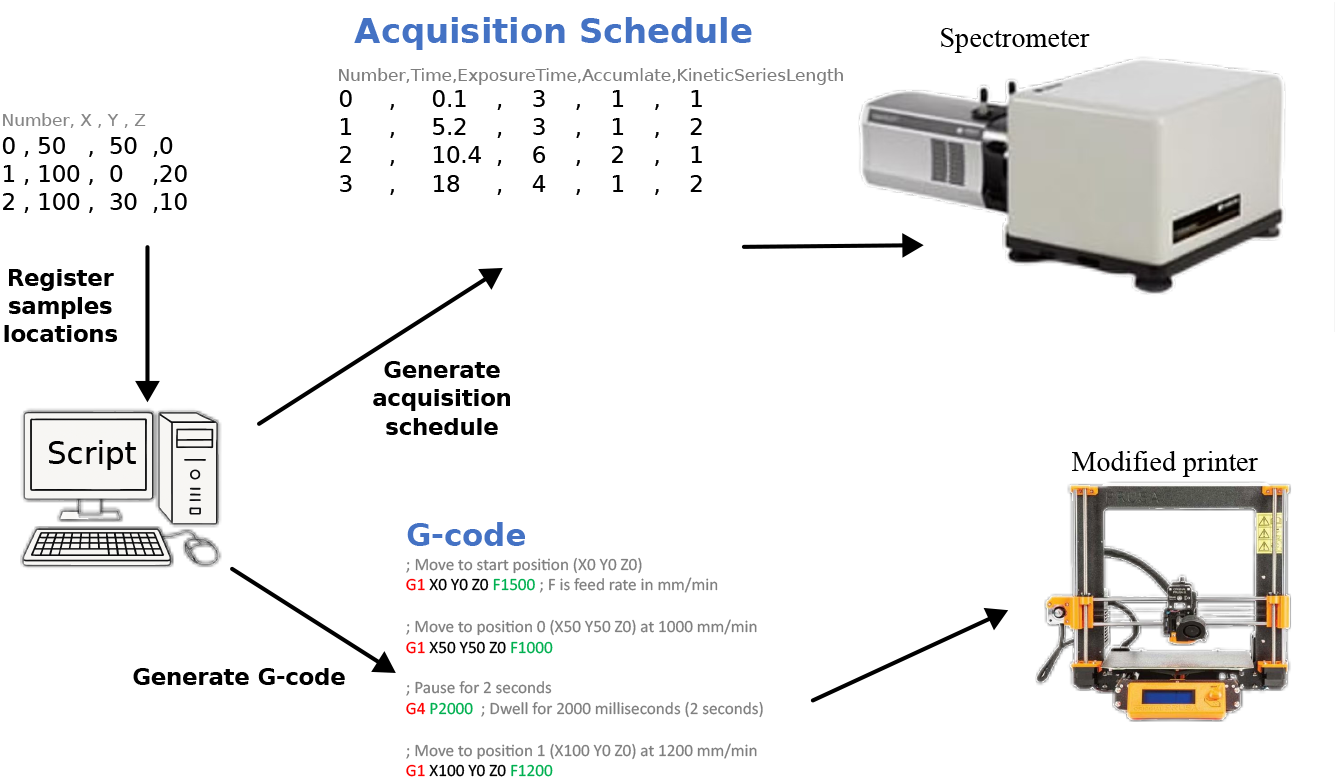
High Throughput Raman. A schematic diagram of the overall workflow of the RamanBot.

G-code offers a variety of motion commands [21]. Among them, the positioning mode commands G90 and G91 for relative and absolute positioning, respectively, are well suited for our device. The relative positioning command G90 is useful when it is required to manually optimize the position of the Raman head for the strongest signal at the first acquisition point and then move relative to this optimized start point. The absolute positioning command G91 is useful when the head is required to start moving from the home point (0,0,0). Furthermore, the G1 motion mode command is used for precise movement.

## Results and Discussion

### The positioning error

Positioning errors in 3D printers originate from several sources, including inaccuracies in the stepper motors controlling motion along the x, y, and z axes, as well as improper belt tension. Excessive belt tension can cause the motor to skip steps, while loose belts may introduce oscillations. Additionally, higher movement speeds increase positioning errors because of the momentum of moving parts, such as the printer bed or the Raman head, resisting rapid changes in motion.

Similarly to the calibration process of a 3D printer, we distinguish positioning errors into two categories: errors along the z-axis, controlled by a single stepper motor, and errors in the xy-plane, controlled by two stepper motors. Errors in the z-direction primarily cause misfocus on the sample, leading to losses in the collected Raman signal. Conversely, imprecision in the xy-plane movement leads to collecting signal form incorrect spatial locations. For these reasons, after assembling the Raman head, which now has a weight different from the original extrusion head, it is critical to evaluate the positioning accuracy. These errors can be compensated for through appropriate G-code commands, if needed.

Before error measurement, the printer bed was carefully leveled to ensure a uniform distance from the Raman head across the entire bed surface as described in [35].

To investigate the positioning error in the Z-axis, Raman signals were collected from ten PLA columns printed with heights ranging from 0.5 to 5 cm, as illustrated in Fig. 3:A. To evaluate the XY-plane error, Raman signals were collected at 16 locations on a 4×4 PLA grid, where the hemispheres are placed at the grid intersections as shown in Fig. 3:B. The dimensions of both tools were measured precisely to minimize geometric errors, and a 100% material fill was used in the printing process to ensure uniform surface material. The test models used in this assessment can be found in [31].

**Fig 3.**
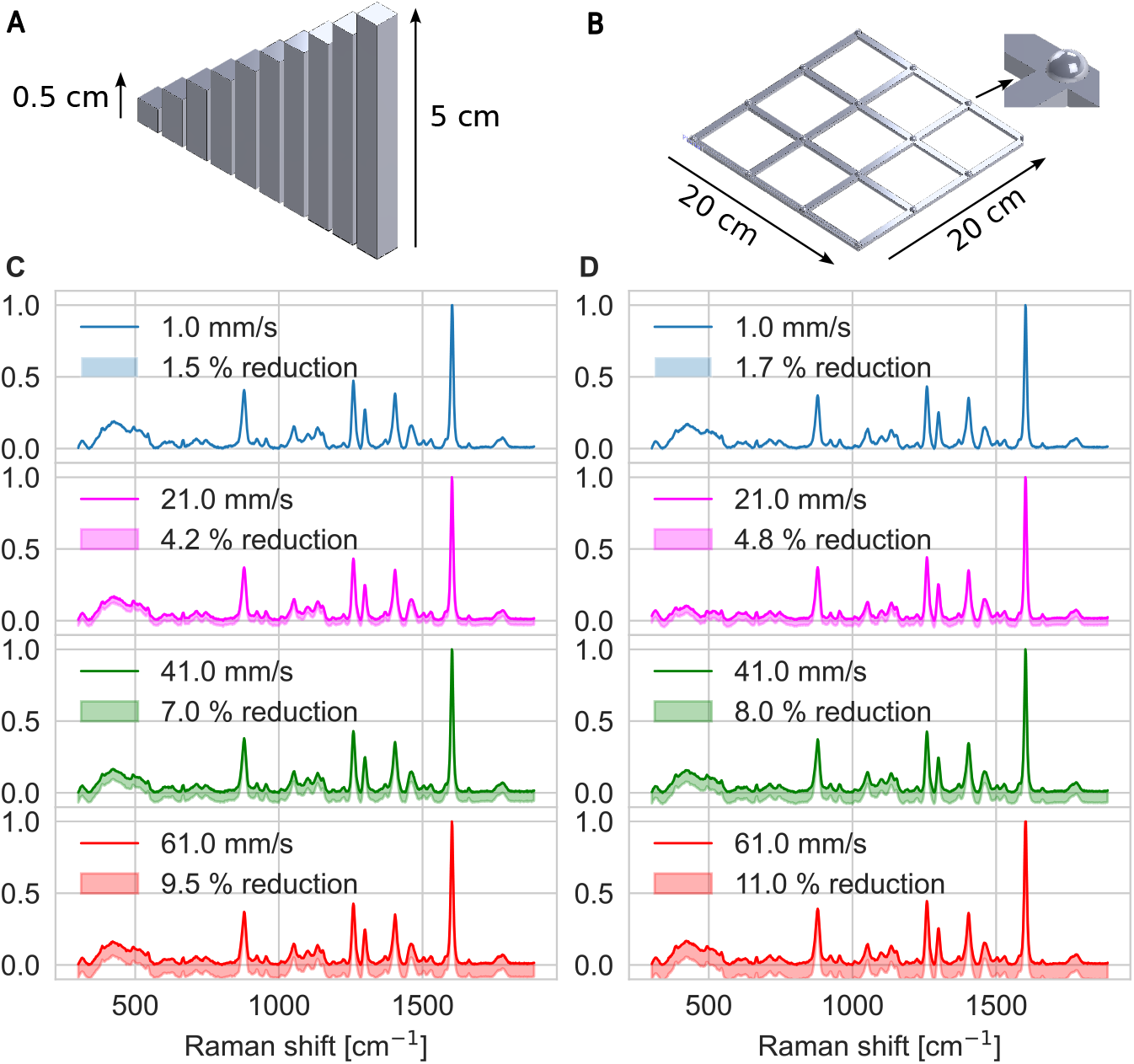
Positioning error. The variation in PLA spectrum due to the mispositioning caused by different speeds in A: Z movements B: XY movements

In order to measure the Z errors, we first calibrated the initial position of the objective lens to be at the focal point of the printing bed. Using the G91 command for relative positioning, we then instructed the machine to move upward by 0.5 mm at a fixed speed. We then placed a 0.5 mm column beneath the Raman head, ensuring its top was at the focal distance from the lens. Any decrease in the recorded Raman intensity at this point would indicate a positioning error. This procedure was then repeated by systematically raising the head in 0.5 mm increments and inserting a corresponding column of increasing height. The experiment was carried out at four different speeds: 1, 21, 41, and 61 mm/s. Figure 3:C shows the Raman signals collected at these speeds, with shaded areas representing how much the signal can drop because of the positioning error. As expected, the slowest speed (1 mm/s) yielded the smallest reduction ( 1.5%). Increasing the speed results in greater focus errors and a corresponding reduction in Raman intensity. The maximum reduction measure was ( 9.5%) recorded at 61 mm/s.

The XY positioning error was assessed by scanning a 4x4 grid of 16 hemispheres, as depicted in Fig. 3:B. First, the maximum Raman signal was recorded at the initial hemisphere. The instrument was then commanded to move to the next hemisphere at a fixed speed, where another Raman signal was collected. A decrease in signal intensity indicated that the measurement head was not precisely positioned at the hemisphere’s apex due to movement errors. The remaining positions were scanned sequentially, row by row, at the same speed. This procedure was repeated at all speeds previously used.

The results, shown in Figure. 3:D, demonstrate a correlation between speed and the reduction in signal intensity, which is directly attributable to positioning imprecision. Consistent with the Z-axis error findings, the slowest speed (1 mm/s) resulted in a minimal signal reduction of approximately 1.7%. In contrast, the highest speed (61 mm/s) caused a much larger reduction of approximately 11%. These findings quantify the reliability of the proposed device at different scanning speeds. It is important to note that the precision of these results is contingent on the quality of the 3D printer components and the accuracy of the system assembly.

### Microplate Plateform

After calibrating the device for optimal performance, we evaluated its capability to read a standard 96-well microplate. Microplates are commonly made of polystyrene, which exhibits a strong Raman peak near 1000 cm^−1^ attributed to the ring breathing mode [36]. Additional, less intense peaks were observed at approximately 1030 cm^−1^ and 1602 cm^−1^.

To ensure that the microplate remains securely positioned during acquisitions, we use the magnetic properties of the build plate by printing the custom plate frame shown in Fig. 4:A with holes designed to house metal inserts. This design enables firm attachment of the plate frame to the build plate. Figure 4:B shows the microplate mounted in the frame and affixed to the build plate.

**Fig 4.**
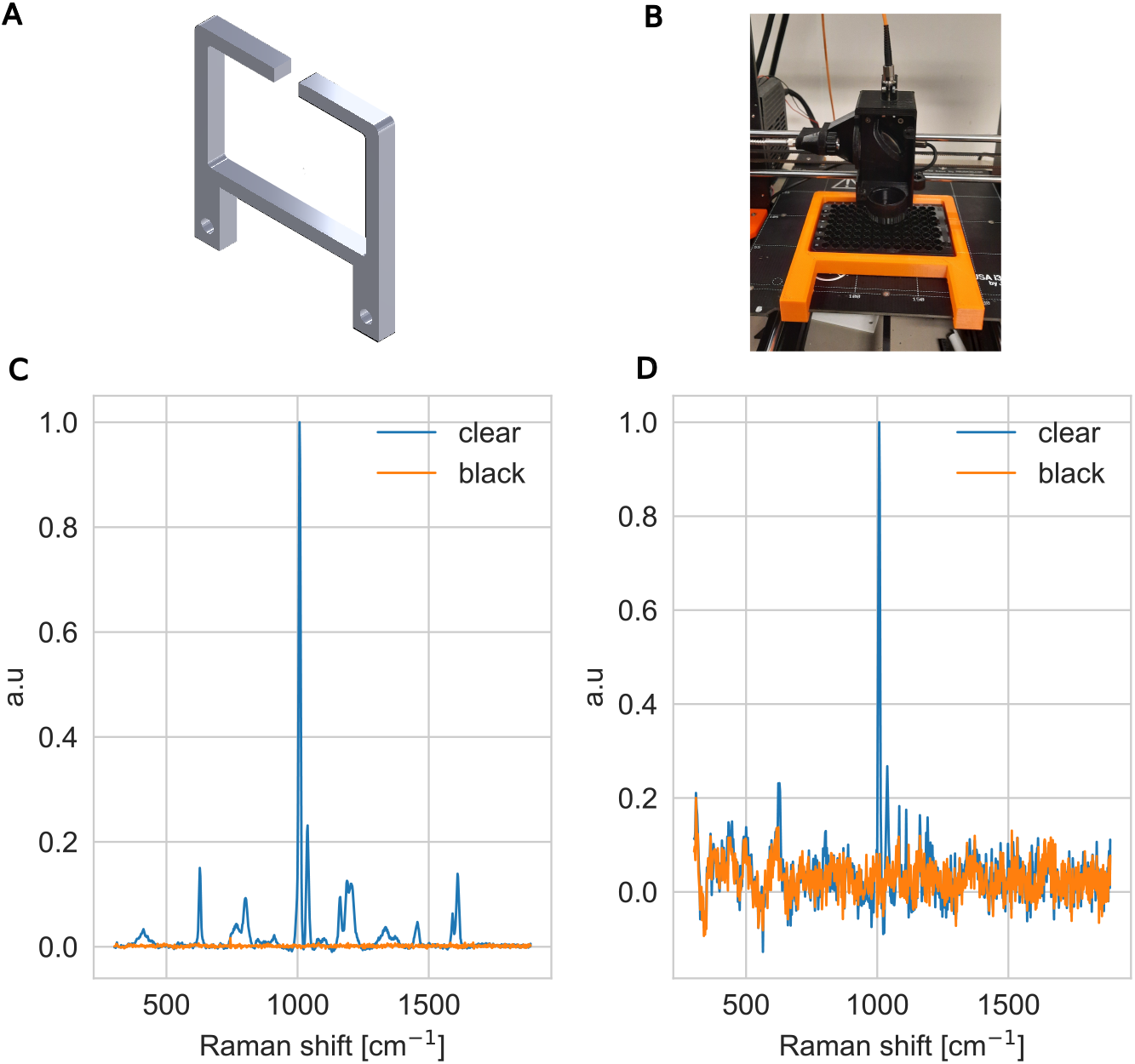
Microplate comparison. The Raman signal captured from clear microplate and a black microplate normalized to the maximum intensity when the well is (c) empty water-filled

Clear microplates are the most commonly used microplates as they allow absorbance assays. However, they cause strong background peaks when used for Raman spectroscopy. In contrast, black microplates produce significantly lower background peaks, as demonstrated in Fig. 4:C. Figure 4:D compares the background Raman peaks of the clear and black microplates when the wells were filled with 330 *µ*L of water. The black microplate exhibits substantially reduced background interference, making it a superior choice for Raman screening applications.

An additional critical factor is the level of cross-talk between adjacent wells. For this evaluation, ethanol was selected because of its strong Raman scattering characteristics. Ethanol has a strong Raman peak around 880 cm^−1^ that corresponds to C-C stretching. Additionally, it has less significant peaks around 1050 cm^−1^,1090 cm^−1^,1280 cm^−1^, and 1455 cm^−1^. These peaks correspond, respectively, to C-O stretching, C-C stretching,C-O-H bending, and CH_2_ / CH_3_ bending. Figure 5 presents Raman spectra collected from a 3×3 well array, where the central well (well 0) was filled with ethanol and the surrounding wells (wells 1 to 8) contain only water. It may be observed that ethanol strong peaks in the surrounding wells were minimal, with only trace amounts of the dominant ethanol peak near 880 cm^−1^ detected in adjacent wells, as highlighted in the inset of Fig. 5. The observed ethanol peaks at well 0 are in agreement with the peaks reported in [37]

**Fig 5.**
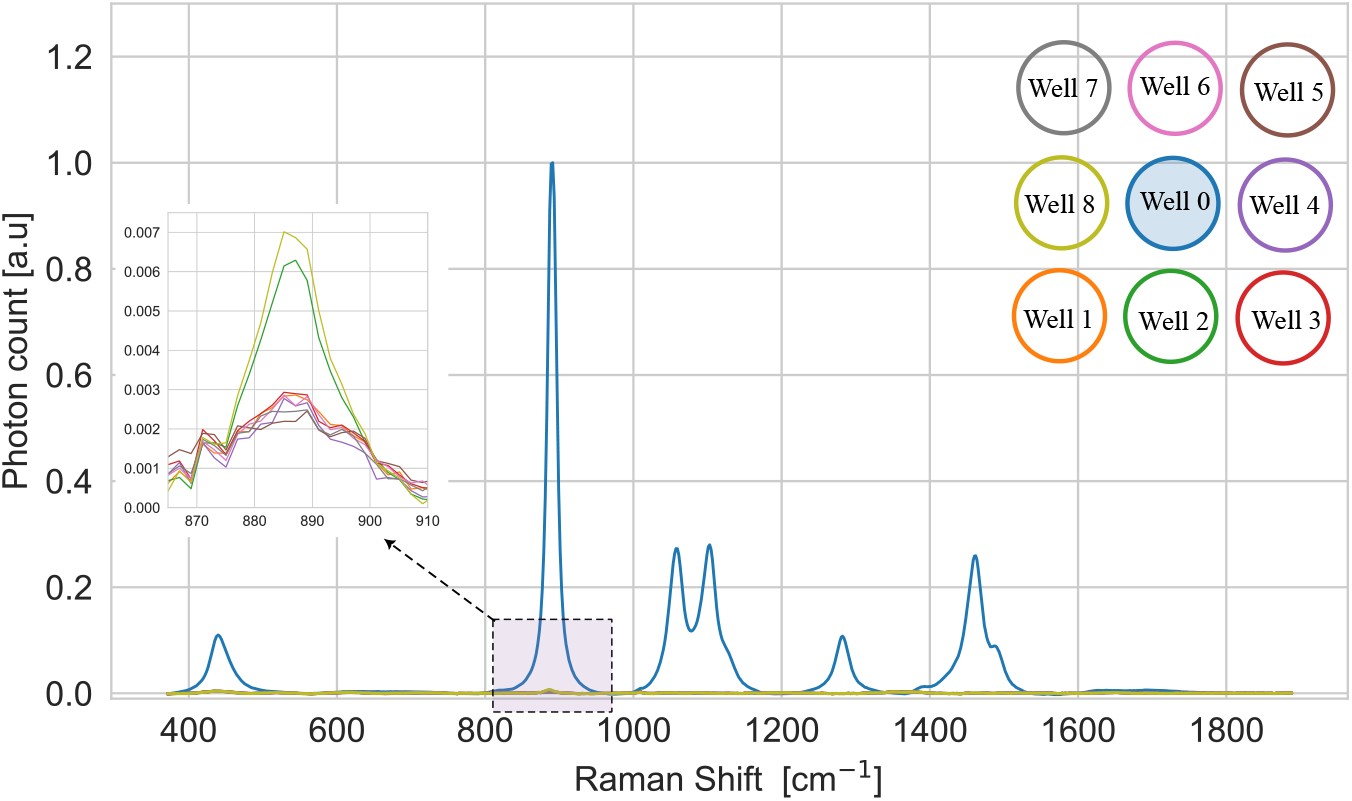
Microplate cross-talk. The Raman signal captured from 3x3 wells with the middle well (well 0) was filled with ethanol and the remaining wells were empty. The inset shows a closer look at the weak traces of ethanol peak at 880 cm^−1^

### Quantitative analysis using 96-well Microplate

In this study, we explore the application of the 96-well microplate for quantitative Raman analysis. Twelve ethanol solutions (columns 1–12) were prepared with concentrations ranging from 200 mM to 2400 mM. Each concentration was replicated eight times (rows A–H), resulting in a total of 96 samples. The samples were loaded into the microplate, which was maintained at a low temperature by placing ice packs on the printing bed to minimize ethanol evaporation during measurements.

The Raman system was programmed to sequentially scan all 96 wells and acquire the Raman spectrum from each. Each well was measured with an acquisition time of 4 seconds. To accommodate system delays and ensure the Raman head fully stabilized before acquisition, a 0.5 second pause was inserted before and after acquisition. The printer speed was set to 20 mm/s, with the home position initialized at well A1. The Raman head was positioned to maximize the signal intensity before scanning was commenced. The total scanning time for the entire microplate was approximately 9 minutes, which is much less than scanning the samples manually while maintaining a low standard deviation between replicas. All collected spectra were processed using the airPLS baseline correction method [38]. Figure 6:A shows Raman spectra of ethanol acquired at different concentrations. A clear increase in Raman intensity is observed with increasing concentration. Figure 6:B shows the average intensity of the dominant ethanol peak at 880 cm^−1^ for the 12 concentrations at each concentration, with error bars indicating the standard deviation. The linear fit yields an R-squared value of 0.9975, which reflects an excellent linear relationship between Raman intensity and ethanol concentration. This is because of the direct relationship between the number of molecules and the intensity of the scattered Raman signal. The maximum recorded standard deviation between replicas was 2.5%, which highlights the reproducibility of the results.

**Fig 6.**
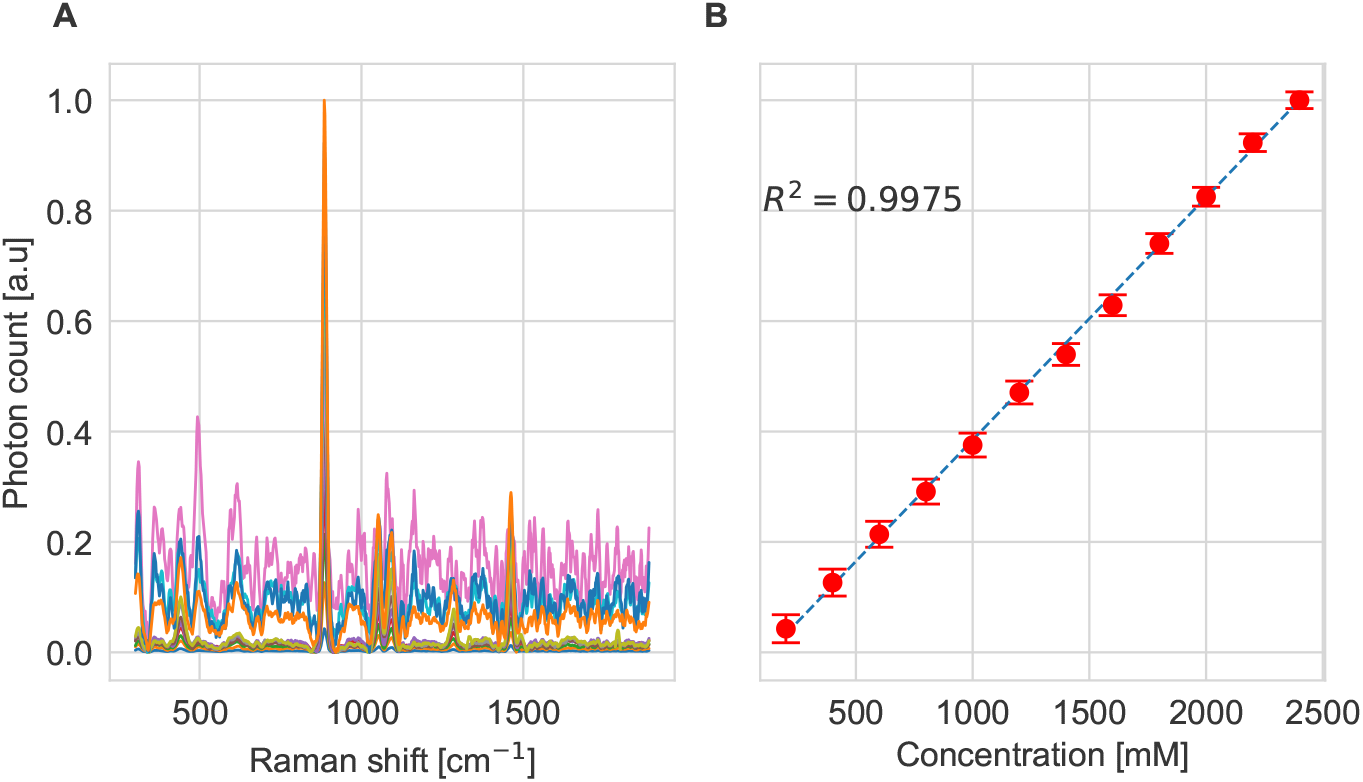
Ethanol concentration. A: The mean of the Raman intensity of different Ethanol concentration normalized to the maximum concentration, B: The Raman intensity at peak 880 cm^−1^ at different concentration fitted to a linear curve.

### Quantitative analysis using microfuge tubes

Microfuge tubes are essential consumables in molecular biology laboratories, designed to withstand rapid temperature changes while their secure sealing minimizes evaporation and contamination. They are widely used in applications such as DNA cloning, gene expression analysis, pathogen detection, and genotyping [39].

To demonstrate the versatility of our proposed Raman system, we designed and 3D printed a holder for the microfuge tubes shown in Fig. 7:A. The holder aligns the samples at an approximate tilt angle of 24°. This inclination allows the Raman head to focus unobstructed on the clear bottom of each tube, maximizing the collected Raman signal. Moreover, the holder is hollow beneath the tubes to prevent interference from Raman signals originating from the PLA material. It is designed to accommodate a 4×3 array of samples, with a spacing of 16 mm between tubes along the x-axis and 45 mm along the y-axis. Two holes were made in the back of the holder to house metal inserts which enable the holder to be magnetically attached to the build plate. Figure 7:B shows the holder mounted on the build plate with the samples in place.

**Fig 7.**
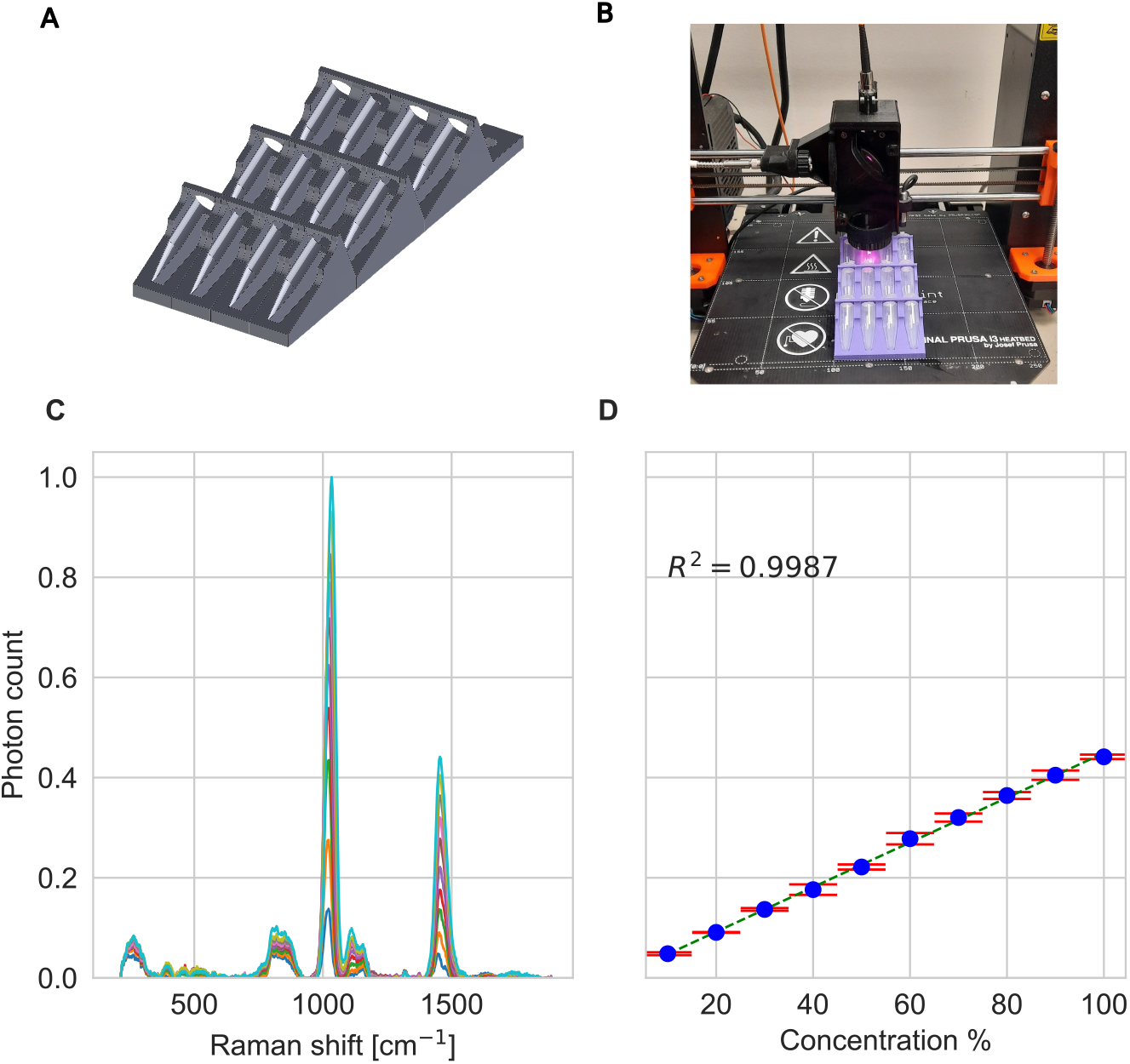
Methanol concentration. Figure shows A: The 3D model of the 4x3 microfuge holder, B: The microfuge holder after being printed and fixed on the build plate, C: The mean of the Raman intensity of different Methanol concentration normalized to the maximum concentration, and D: The Raman intensity at peak 1450 cm^−1^ at different concentration fitted to a linear curve.

We used the holder to analyze 10 methanol concentrations ranging from 10% to 100%. Six different replicas of each concentration were created. The focus was initially optimized on the first sample in the array to maximize Raman signal collection, accounting for the thin microfuge tube wall between the methanol and the excitation laser. Using the known dimensions of the holder, a G-code was generated to sequentially move the Raman head between samples at a speed of 20 mm/s. The acquisition time per sample was set to 20 seconds, with a device pause time of 22 seconds to ensure stabilization before measurement. The total scanning time per replica set was approximately 4.8 minutes. This procedure was repeated for the remaining five replicates. Performing the same experiment manually would take longer time and low reproducibility.

All collected spectra were corrected for the baseline using the airPLS algorithm [38], and the background spectrum of polypropylene was subtracted to isolate the methanol signal. Figure 7:C shows the Raman spectra for each methanol concentration averaged across the replicas. Raman peaks are observed at 1035 cm^−1^, 1450 cm^−1^, and 1125 cm^−1^, which agree well with the peaks reported in [40]. Figure. 7:D tracks the intensity of the methanol peak at 1450 cm^−1^. The results demonstrate a linear relationship between Raman intensity and methanol concentration with error bars representing the standard deviation of the replicates. The linear fit, represented by the dotted line, yields an R-squared value of 0.9987. This indicates excellent quantification performance. The linear behavior is attributed to the linear relationship between the Raman intensity and the number of molecules. The maximum standard deviation noticed was 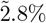, which underscores the reliability of the proposed device.

### System adaptability: Egg-shell Analysis

The programmable flexibility of the RamanBot provides a practical approach to Raman screening in the food industry. It can allow screening to be performed directly within the original packaging of food products. To demonstrate this, we analyzed six eggs placed in their standard 3×2 egg carton, as shown in Fig. 8:A. The egg carton was aligned with the grid lines of the build plate. The center-to-center distance between adjacent eggs was measured to be approximately 45 mm. Accurate knowledge of the relative positions of each egg is essential to enable precise movement between samples. Additionally, the height of each egg was measured to ensure correct focus adjustment for each measurement.

**Fig 8.**
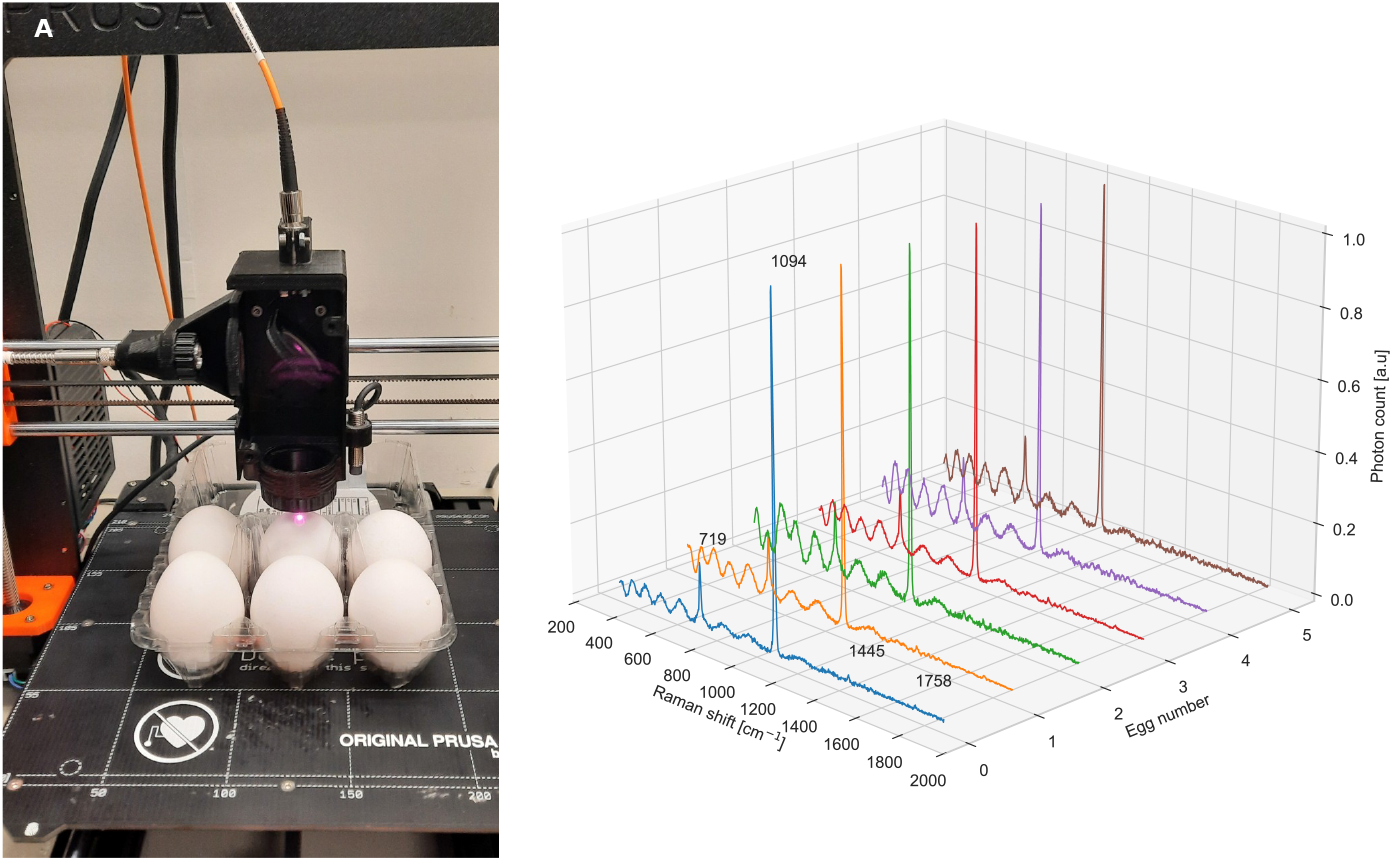
Egg-shell spectrum. Figure shows A: The 3x2 egg package placed under Raman Head, B: The Raman signal captured for all the 6 eggs.

For this experiment, the Raman acquisition time per sample was set to 4 seconds. To account for potential delays on the spectrometer side and to allow sufficient time for data saving and optical stabilization after stopping, the device pause time at each egg was set to 6 seconds. This pause also mitigates any vibrations caused by sudden halts. The printer speed was maintained at 20 mm/s. Using this information, a G-code script was generated to sequentially scan the six samples, and the acquisition schedule was generated to synchronize the system with the spectrometer. The total screening time for this experiment was approximately 95 seconds. This recorded time highlights the time efficiency of the device compared to manual screening. An extension of this idea is to use generated G-code to scan similar egg cartons in the market, reducing both labor and time.

After acquisition, baseline correction was applied to the spectra using the airPLS algorithm [38]. Figure 8:B displays the Raman spectra obtained from the eggshells. A dominant peak is observed at 1085 cm^−1^, indicative of crystalline calcium carbonate (CaCO_3_), consistent with previous reports [41, 42]. Additional bands appeared at 719 cm^−1^, 1445 cm^−1^, and 1758 cm^−1^, corresponding to symmetric deformation of CO_2_, CH_2_ bending, and C-O stretching vibrations, respectively, in agreement with literature [41]. The sinusoidal pattern observed between 300 cm^−1^ and 1000 cm^−1^ is attributed to interference effects from the eggshell’s multilayer structure.

## Conclusion

We have presented a high-throughput Raman screening system developed by modifying the XYZ motion stage of a standard 3D printer. The central innovation lies in the design of a custom Raman head that replaces the extrusion head and integrates all optical components necessary for sample excitation and Raman signal collection. This approach enables flexible, high-throughput screening and holds potential to be extended to other spectroscopic modalities. Moreover, it opens the possibility of incorporating Raman spectrometers into existing 3D printing platforms. The low cost of the proposed system makes Raman measurements more accessible to laboratories.

The complete system workflow has been detailed, including G-code generation and acquisition scheduling. We have also introduced a method to evaluate the effect of positioning errors on the Raman signal intensity. The speed and reproducibility of the system were demonstrated through quantitative analyses of ethanol in a microplate. Additionally, the feasibility of fabricating a microfuge tube array for quantitative methanol analysis has been presented, which highlights the versatility of the device. Finally, the adaptability of the system to unconventional sample arrangements was demonstrated by successfully acquiring Raman spectra from six eggs within their carton.

## Notes

### Competing Interest Statement

The authors have declared no competing interest.

## References

1. Pelletier MJ. Quantitative Analysis Using Raman Spectrometry. Applied Spectroscopy. 2003;57(1). doi:10.1366/000370203321165133.

2. Taylor LS, Zografi G. The quantitative analysis of crystallinity using FT-Raman spectroscopy. Pharmaceutical Research. 1998;15(5):755–761. doi:10.1023/a:1011979221685.

3. Shipp DW, Sinjab F, Notingher I. Raman spectroscopy: techniques and applications in the life sciences. Advances in Optics and Photonics. 2017;9(2). doi:10.1364/aop.9.000315.

4. Stone N, Kendall C, Shepherd N, Crow P, Barr H. Near-infrared Raman spectroscopy for the classification of epithelial pre-cancers and cancers. Journal of Raman Spectroscopy. 2002;33(7). doi:10.1002/jrs.882.

5. Kendall C, Stone N, Shepherd N, Geboes K, Warren B, Bennett R, et al. Raman spectroscopy, a potential tool for the objective identification and classification of neoplasia in Barrett’s oesophagus. The Journal of Pathology. 2003;200(5). doi:10.1002/path.1376.

6. Macarron R, Banks MN, Bojanic D, Burns DJ, Cirovic DA, Garyantes T, et al. Impact of high-throughput screening in biomedical research. Nature Reviews Drug Discovery. 2011;10(3). doi:10.1038/nrd3368.

7. Jenkins CA, Jenkins RA, Pryse MM, Welsby KA, Jitsumura M, Thornton CA, et al. A high-throughput serum Raman spectroscopy platform and methodology for colorectal cancer diagnostics. The Analyst. 2018;143(24):6014–6024. doi:10.1039/c8an01323c.

8. Schie IW, R”uger J, Mondol AS, Ramoji A, Neugebauer U, Krafft C, et al. High-throughput screening Raman spectroscopy platform for label-free cellomics. Analytical chemistry. 2018;90(3):2023–2030.

9. Liao HX, Bando K, Li M, Fujita K. Multifocal Raman spectrophotometer for examining drug-induced and chemical-induced cellular changes in 3D cell spheroids. Analytical Chemistry. 2023;95(39):14616–14623.

10. Seo BW, A. Ja’farawy MS, Jung HS, Choi YW, Park SG, Choi WJ. Miniaturized and Automated Optical-Switch Raman Spectroscopy Enabling Multiwell Surface-Enhanced Raman Spectroscopy Screening More than 26,000 Wells per Day. BioChip Journal. 2025; p. 1–11.

11. Zhou B, Qu C, Du S, Gao W, Zhang Y, Ding Y, et al. Multi-analyte High-Throughput Microplate-SERS Reader with Controllable Liquid Interfacial Arrays. Analytical Chemistry. 2022;94(21). doi:10.1021/acs.analchem.2c00252.

12. Medipally DKR, Maguire A, Bryant J, Armstrong J, Dunne M, Finn M, et al. Development of a high throughput (HT) Raman spectroscopy method for rapid screening of liquid blood plasma from prostate cancer patients. The Analyst. 2017;142(8):1216–1226. doi:10.1039/c6an02100j.

13. Kawagoe H, Ando J, Asanuma M, Dodo K, Miyano T, Ueda H, et al. Multiwell Raman plate reader for high-throughput biochemical screening. Scientific Reports. 2021;11(1). doi:10.1038/s41598-021-95139-8.

14. Wolf S, Popp J, Frosch T. Multifocal hyperspectral Raman imaging setup for multi-well plates. Sensors and Actuators B: Chemical. 2023;375:132949.

15. Heid CA, Stevens J, Livak KJ, Williams PM. Real time quantitative PCR. Genome Research. 1996;6(10). doi:10.1101/gr.6.10.986.

16. Tripathy S, Chavva S, Cote GL, Mabbott S. Modular and handheld Raman systems for SERS-based point-of-care diagnostics. Current Opinion in Biomedical Engineering. 2023;28:100488.

17. Hargreaves M. Handheld Raman, SERS, and SORS. Portable spectroscopy and spectrometry. 2021; p. 347–376.

18. Beganović A, Hawthorne LM, Bach K, Huck CW. Critical review on the utilization of handheld and portable Raman spectrometry in meat science. Foods. 2019;8(2):49.

19. Kranenburg RF, Verduin J, de Ridder R, Weesepoel Y, Alewijn M, Heerschop M, et al. Performance evaluation of handheld Raman spectroscopy for cocaine detection in forensic case samples. Drug testing and analysis. 2021;13(5):1054–1067.

20. Oberg E, Jones FD, Horton HL, Ryffel HH, Mccauley CJ, Heald R, et al. Machinery’s handbook. vol. 6. Industrial Press New York; 1914.

21. Gcode;. Available from: https://marlinfw.org/meta/gcode/.

22. Gibson I, Rosen D, Stucker B, Khorasani M. Additive Manufacturing Technologies. Springer International Publishing; 2021. Available from: 10.1007/978-3-030-56127-7.

23. Ivanova TN, Bialy W, Nordin V. Improvement of grinding technology with vortex cooling of steels that are liable to crack propagation. Multidisciplinary Aspects of Production Engineering. 2019;2.

24. Hooper S. CoreXY Kinematics; 2021. 3D Distributed. Available from: https://www.3ddistributed.com/corexy-kinematics/.

25. Tay SH, Choong WH, Yoong HP. A review of scara robot control system. In: 2022 IEEE International Conference on Artificial Intelligence in Engineering and Technology (IICAIET). IEEE; 2022. p. 1–6.

26. Arawade SS. State of Art Review on SCARA Robotic Arm. International Journal of Advanced Research in Science, Communication and Technology. 2021;3(1):160–166. All3DP. The Types of FDM 3D Printers: Cartesian, CoreXY & More; 2025.

27. All3DP. Available from: https://all3dp.com/2/cartesian-3d-printer-delta-scara-belt-corexy-polar/.

28. Elghitany MN, Ahmed A, Zaki D, Behhit D, Hosni H, Nour H, et al. Advancements in Design, Kinematics, and Control: A Comprehensive Review of Delta Robot Research. Advanced Sciences and Technology Journal. 2024;1(2):1–38.

29. Muniyandi GB. In-Depth Analysis of Kinematic, Dynamic, and Control Aspects of a 4-Axis SCARA Robot Manipulator. International Journal of Robotic Engineering. 2024;7(1).

30. GitHub - prusa3d/Original-Prusa-i3: Original Prusa i3 MK2 3D printer printed parts;. https://github.com/prusa3d/Original-Prusa-i3. Available from: https://github.com/prusa3d/Original-Prusa-i3.

31. Atia K. RamanBot; 2025. https://github.com/khaledAtia/RamanBot.

32. Andor Technology. HoloSpec On-axis high throughput imaging spectrograph; 2025. https://andor.oxinst.com/assets/uploads/products/andor/documents/Andor-Holospec-Specifications.pdf.

33. Kliment. Pronterface: Graphical host software from the Printrun 3D-printing suite; 2014. Available from: http://www.pronterface.com/.

34. Oxford Instruments Andor Ltd. Solis Software; 2025. https://andor.oxinst.com/products/solis-software/.

35. Prusa Research. Bed Level Correction; 2025. https://help.prusa3d.com/article/bed-level-correction_2267.

36. Vandenabeele P, Edwards HGM, Moens L. A Decade of Raman Spectroscopy in Art and Archaeology. Applied Spectroscopy. 2000;54(12):1553–1563. doi:10.1366/0003702001950481.

37. Burikov S, Dolenko T, Patsaeva S, Starokurov Y, Yuzhakov V. Raman and IR spectroscopy research on hydrogen bonding in water–ethanol systems. Molecular physics. 2010;108(18):2427–2436.

38. Zhang ZM, Chen S, Liang YZ. Baseline correction using adaptive iteratively reweighted penalized least squares. The Analyst. 2010;135(5):1138. doi:10.1039/b922045c.

39. Ausubel FM, Brent R, Kingston RE, Moore DD, Seidman JG, Smith JA, et al. Current Protocols in Molecular Biology. Wiley; 1994.

40. Vaskova H. Spectroscopic determination of methanol content in alcoholic drinks. Int J Biol Biomed Eng. 2014;8:27–34.

41. Ferraz E, Gamelas JAF, Coroado J, Monteiro C, Rocha F. Eggshell waste to produce building lime: calcium oxide reactivity, industrial, environmental and economic implications. Materials and Structures. 2018;51(5). doi:10.1617/s11527-018-1243-7.

42. Prabakaran K, Balamurugan A, Rajeswari S. Development of calcium phosphate based apatite from hen’s eggshell. Bulletin of Materials Science. 2005;28(2). doi:10.1007/bf02704229.

